# SARS-CoV-2 neutralization and protection of hamsters via nasal administration of a humanized neutralizing antibody

**DOI:** 10.1101/2024.07.08.602557

**Authors:** Mikhail Lebedin, Nikolai Petrovsky, Kairat Tabynov, Kaissar Tabynov, Yuri Lebedin

## Abstract

Monoclonal antibodies are widely used for the treatment of infectious human diseases, including COVID-19. Since the start of the pandemic, eight monoclonal antibodies against SARS-CoV-2 were granted emergency use authorization. High mutation rate of the SARS-CoV-2 virus has led to the emergence of highly transmissible variants efficiently evading vaccine-induced immunity. This highlights the importance of identifying broadly neutralizing antibodies with therapeutic potential. In this study, we used a panel of murine monoclonal antibodies (mAb) to identify a subset that bound and neutralized a broad spectrum of SARS-CoV-2 variants. Intranasal delivery of XR10, the most promising murine mAb, protected hamsters against infection by Alpha and Delta variants. We next humanized XR10 mAb using a combination of CDR-grafting and Vernier zones preservation approaches (CRVZ) to create a panel of humanized antibody variants. We ranked the variants based on their spike binding ability and virus neutralization. Of these, XR10v48 demonstrated the best ability to neutralize SARS-CoV-2 variants. XR10v48 was protective in hamsters when given as a single 50 µg/kg intranasal dose at the time of viral challenge. XR10v48 features 34 key amino acid residues retained from the murine progenitor. Our work introduces a potent humanized antibody that demonstrates neutralizing activity *in vivo* at a low dose.

## INTRODUCTION

Since the first outbreak in December 2019 [1], World Health Organization has reported 775.7 million cases of coronavirus disease 2019 (COVID-19) with the death toll reaching 7.0 million globally by July 2024 (https://data.who.int/dashboards/covid19). The causal agent, a beta coronavirus referred to as severe acute respiratory syndrome virus 2 (SARS-CoV-2) [2], bears a spike protein crucial for binding the host receptor, angiotensin-converting enzyme 2 (ACE2) [3]. Host receptor-spike interaction is mediated by a domain denoted as the receptor-binding domain (RBD) [4], which includes the receptor-binding motif (RBM), a major target of neutralizing antibodies [5]. SARS-CoV-2 demonstrates a high mutation rate resulting in several variants of concern reported since the beginning of the pandemic, including Alpha (B.1.1.7), Beta (B.1.351), Gamma (P.1), Delta (B.617.2), Omicron (B.1.1.529), featuring higher severity of disease (Delta, [6]) and transmissibility (Omicron [7]). Most mutations in the currently predominant Omicron variant are located in the spike RBD, which contributes to the virus immune evasion by reducing or completely abrogating the binding of neutralizing antibodies [8] thereby diminishing vaccination efficiency [9–11].

Since hybridoma technology was introduced in 1975 [12], monoclonal antibodies (mAbs) generated through mouse immunization have become a valuable tool in basic research and therapy. In 1986, the first murine mAb Muronomab-CD3 was approved by the United States Food and Drug Administration (FDA) to control kidney transplant rejection [13]. To date, 130 mAbs are approved for the treatment of various types of cancer, autoimmune disorders, and viral infections [14–17]. So far, eight mAbs neutralizing SARS-CoV-2 were granted emergency use authorization (EUA): bamlanivimab, etesivimab, casirivimab, imdevimab, sotrovimab, cilgavimab, tixagevimab, bebtelovimab, with only the latter being effective in Omicron BA.1 cases [18].

Clinical application of mouse mAbs is rendered inefficient by human anti-mouse antibodies(HAMA), which can lead to therapeutic agent clearance as seen in about 50% of patients receiving murine antibodies [19,20]. Advances in genetic engineering and recombinant technology yielded several techniques that seek to overcome the HAMA response, including transgenic mice expressing human IgM, IgG, and IgK [21], human antibody phage display [22], and murine mAb humanization [23]. Humanization aims to increase the similarity of the administered antibody to human immunoglobulins, minimizing anti-drug reactivity and extending its half-life in the organism while retaining its antigen-binding properties [24]. Several methods have been developed to achieve mAb humanization spanning from constant domain replacement [25] to antibody resurfacing [26] or specificity-determining residues grafting [27]. Constant domain replacement was successfully used to reduce the immunogenicity of several FDA-approved chimeric antibodies including abciximab [25], rituximab [14], and cetuximab [28]. Chimeric mAb preserve murine content in the variable domain, presenting a potential target for HAMA response. The risk of the anti-drug response can be further decreased by complementarity-determining region grafting (CDR grafting) in which the murine CDRs are transplanted onto the human frameworks (FRs) [29]. In most cases, direct grafting of the CDR loops onto human FRs leads to substantial loss of target affinity [30]. Several buried residues comprising the β-sheet framework regions, referred to as Vernier zones residues (VZRs), were reported to influence the conformation of the CDR loops, therefore affecting the antigen binding [31,32]. Therefore, CDR grafting should be accompanied by transferring the corresponding VZRs while controlling the affinity of the humanized antibody.

Here we report the generation of a large panel of murine mAbs, a subset of which potently neutralized a range of SARS-CoV-2 variants. The lead candidate, XR10, was protective when delivered nasally in a hamster challenge model and was then humanized by the CRVZ approach. We selected a humanized variant with the highest neutralizing activity and demonstrated its effectiveness against the SARS-CoV-2 Delta strain in a hamster challenge model.

## RESULTS

### Generation of murine SARS-CoV-2 neutralizing antibodies

To generate the mAbs, we immunized BALB/c mice with Wuhan SARS-CoV-2 Spike protein and used hybridoma technology to create the clones (Figure 1A). Purified mAbs were tested for binding to Spike and RBD proteins from various virus strains and virus neutralization (Figure 1B). XR10, XR14, and XR51 mAbs featured high binding to various Spike proteins, >90% inhibition of ACE2-RBD binding, and highest neutralizing activity (IC_50_ = 17.7, 28.8, 12.2 ng/ml for XR10, XR14, and XR51, accordingly) toward the Wuhan variant and were selected for humanization. While XR14 was not able to neutralize Delta or Omicron BA.1 variants, XR10 showed moderate neutralization capacity against Omicron BA.1 (410 ng/ml) and XR51 neutralized the Delta variant with IC_50_ 65.5 ng/ml (Figure 1B, Table S1).

**Figure 1.**
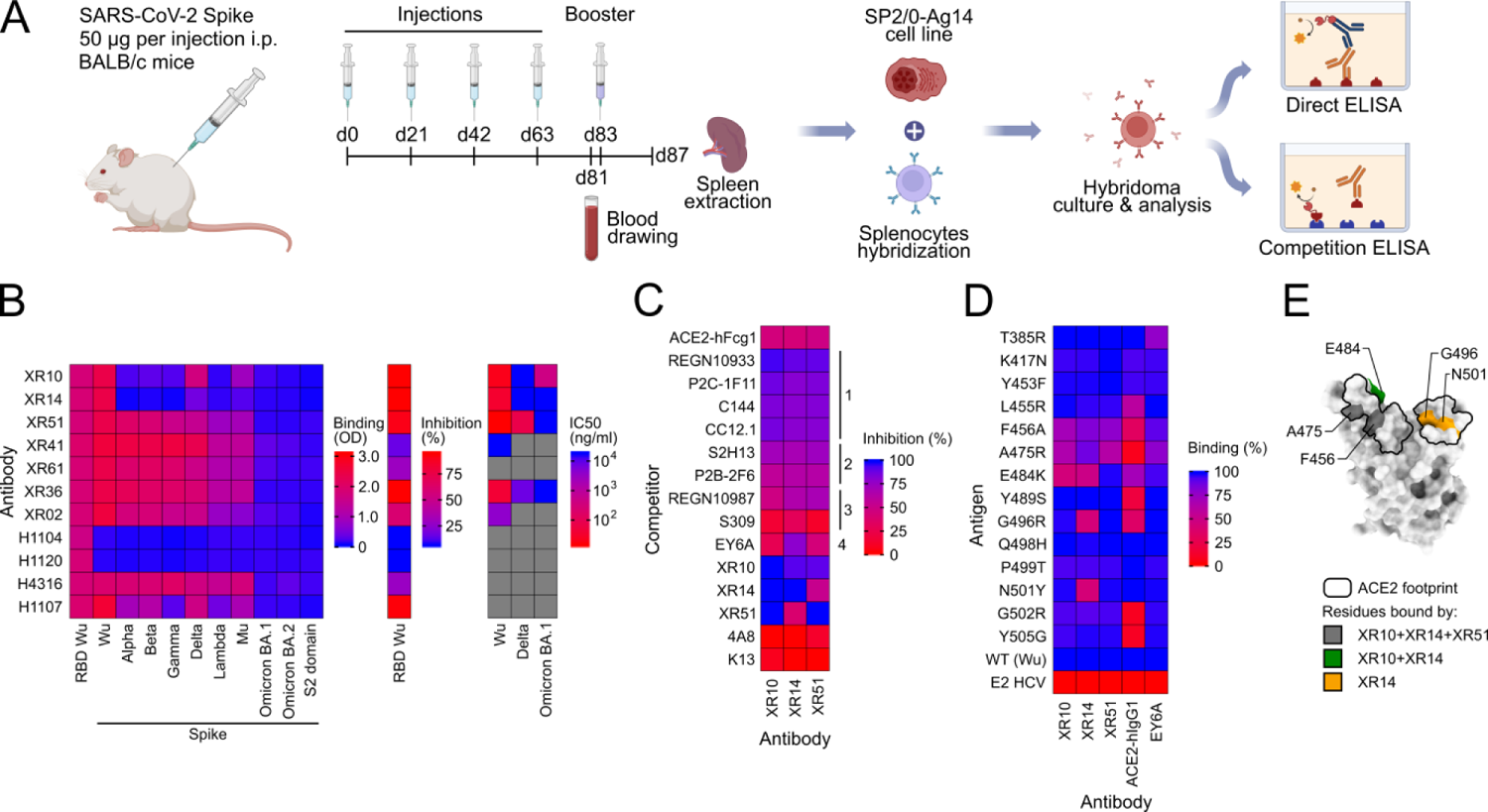
A set of mAbs against SARS-CoV-2 Spike efficiently bind and neutralize various viral strains. **(A)** Scheme depicting mice immunization, hybridoma generation, and mAb analysis. **(B)** mAbs were screened for binding to a diverse set of SARS-CoV-2 Spike proteins (on the left), for inhibition of ACE2-Wuhan RBD binding (middle), and neutralization of the live viruses (on the right). H1104, H1120, H4316, H1107 – commercial antibody controls. **(C)** Wuhan RBD binding competition with various antibodies interacting with distinct epitopes on the RBD surface. K13 - anti-human CD81 antibody used as a negative control. **(D)** mAb binding to a set of Wuhan RBD mutants. E2 - hepatitis C virus surface protein subunit used as a negative control. **(E)** Residues affecting the XR mAbs binding and the ACE2 footprint were mapped on the Wuhan RBD surface.

### mAbs epitope characterization

To map the epitope targeted by the selected mAbs *in vitro*, we performed a binding competition assay with ACE2 and a panel of previously characterized antibodies [33–41] (Figure 1C, Table S1). We used the antibodies representative of different classes: class 1 defined by targeting the RBM in “up” conformation; class 2 that bind the RBM in both “up” and “down” conformations; class 3 that bind outside of ACE2-interacting interface but are able to hinder the host receptor interaction; and class 4 that do not perturb the ACE2-RBD binding [42]. All three mAbs competed moderately with ACE2-hFcg1 binding (44.0%, 42.4%, 47.7% inhibition for XR10, XR15, and XR51) and demonstrate >80% binding inhibition by such class 1 antibodies as REGN10933 (90.4-95.9%), P2C-1F11 (81.5-88.4%), C144 (80.8-84.5%), and CC12.1 (80.8-86.7%). All selected antibodies competed moderately with class 2 antibodies, as P2B-2F6 (56.6-62.9%) and S2H13 (69.0-73.6%) and competed weakly with S309, which is classified as a epitope 3 antibody (9.9-28.5%). Surprisingly, XR14 showed 79.8% binding inhibition for the class 4 antibody EY6A. Cross-inhibition assay shows that XR10 epitope overlapped with both XR14 and XR51 interaction zones (93.1% and 91.8% inhibition, correspondingly), while we detected a weak inhibition for XR51-XR14 binding (38.2%). These data suggest that XR10 and XR51 bind an RBM epitope and are closely related to class 1/2 Abs, while XR14 might demonstrate a unique multimodal binding overlapping with class 1/2 and class 4 Abs.

To define the epitopes on the amino acid residue level, we performed binding assay with various Wuhan RBD mutants. Mutations F456A, A475R, E484K, located within class 1/2 epitopes, substantially decreased the binding of XR10 and XR14 mAbs, while XR51 bound to E484K mutant with nominal affinity (Figure 1D, Table S1). While not affecting XR10 and XR51, mutations G496S and N501Y, located in the right shoulder of RBD (Figure 1E) [43], reduced the XR14 binding to 43.1% and 42.2%, accordingly. These data are in agreement with competition analysis and indicate that all three antibodies belong to class 1 and class 2 antibodies. Furthermore, the XR10 epitope overlapped with both XR14 and XR51 interaction zones, while XR14 and XR51 featured distinct modes of binding.

### XR10, XR14, and XR51 humanization

To humanize the selected mouse antibodies, we identified the human germline (HGL) heavy and light chain variable domain sequences of the highest homology by aligning the murine AA V(D)J sequences to the library of human segments using IgBLAST (Figure S1). We assumed no junctional diversity (no P/N-nucleotides addition) when reconstructing HGL sequences. We introduced the humanization index (DtH, distance to human, indicating the total number of AA exchanges needed to be introduced in heavy and light variable domains to arrive to the HGL sequence) to reflect the depth of humanization. Mouse XR10, XR14, and XR51 display DtH of 57, 69, and 50 AAs exchanges, correspondingly (Figure S1). Taken that the total length of heavy and light chain variable domains are 231, 226, and 235 AAs for XR10, XR14, and XR51, it translates to 75.3%, 69.5%, and 78.7% homology to HGL, respectively. CDRs predicted by IgBLAST software were grafted into fully human frameworks and constant regions were replaced with human IgG1 or IgK (CG version, Table S2). Additionaly, we attempted to replace the amino acid residues located on the flanks of CDR loops to create deeply humanized mAb variants (DH, Table S2). CDR grafting decreased DtH from 57, 69, 50 to 24, 19, 12 (89.6%, 91.6%, 94.9% homology to HGL) for XR10, XR14, and XR51, correspondingly, while deep humanization minimized the DtH to 12, 11, 3 (94.8%, 95.1%, 98.7% homology to HGL). Binding analysis revealed a drastic drop in affinity for humanized antibodies with only CG-XR10 binding to Wuhan RBD with <1.0 ng/µl EC_50_ (Figure 2A, Table S1). These results indicate a significant contribution of Vernier zones residues (VZRs) to antigen recognition. To locate the critical VZRs we assessed the contribution of the heavy and the light chains by generating all combinations of the murine, CG, and DH variants (Figure 2B). For XR10 and XR14 the binding is governed by both chains, while for XR51, the nominal binding is retained if the original murine heavy chain is coupled with any light chain variant, pointing to virtually exclusive heavy chain-mediated binding (Figure 2B, Table S1). To further narrow down the location of the critical VZRs, we performed framework reshuffling for both chains of XR10 and XR14 (Figure 2C,D, Table S1) and for heavy chain of XR51 (Figure 2E, Table S1). Surprisingly, XR51 framework reshuffling revealed all-or-nothing effect of heavy chain FR2 and FR3 mutagenesis, i.e. the mAb binds Wuhan RBD with original affinity only if both HC FR2 and FR3 are not humanized. This allowed us to isolate the critical VZRs for XR51 heavy chain. While A50 is immediately adjacent to CDR2, K67, A68, L70, A72, and S76 are located in the middle of FR3 with CDR2-proximal FR3 residues (S59, N61, R63, K65) being insensitive to humanization. These data suggest that Vernier zone residues shall be identified experimentally as they might be located in buried sites outside of interaction zone. We summarized an experiment-driven humanization pipeline by CDR-grafting and Vernier zone preservation approach (CRVZ) in Figure S2. Of note, Humanization yielded XR10v43 and XR51v127 variants with high level of humanization and nominal or higher-than-nominal binding strength (XR10v43: DtH=27, 88.3% homology to HGL, EC_50_=0.016 ng/µl; XR51v127: DtH=30, 87.2% homology to HGL, EC_50_=0.006 ng/µl, Figure 2F, Table S1). We compared our humanization pipeline to previously published machine-learning approach called Hu-mAb [44]. Predicted humanization mutations mostly overlap with the set introduced in this study, although Hu-mAb uses different germline segments for XR10 and XR51 kappa chains (Figure S1). Humanness score estimated by Hu-mAb strongly correlated to DtH (r = −0.91) while offering more discrete values (Figure S3A). Binding strength demonstrated similar dependence from humanness score in comparison to DtH (Figure S3B). Therefore, both approaches can be used to assess the extent of humanization and design a humanized mAb sequence.

**Figure 2.**
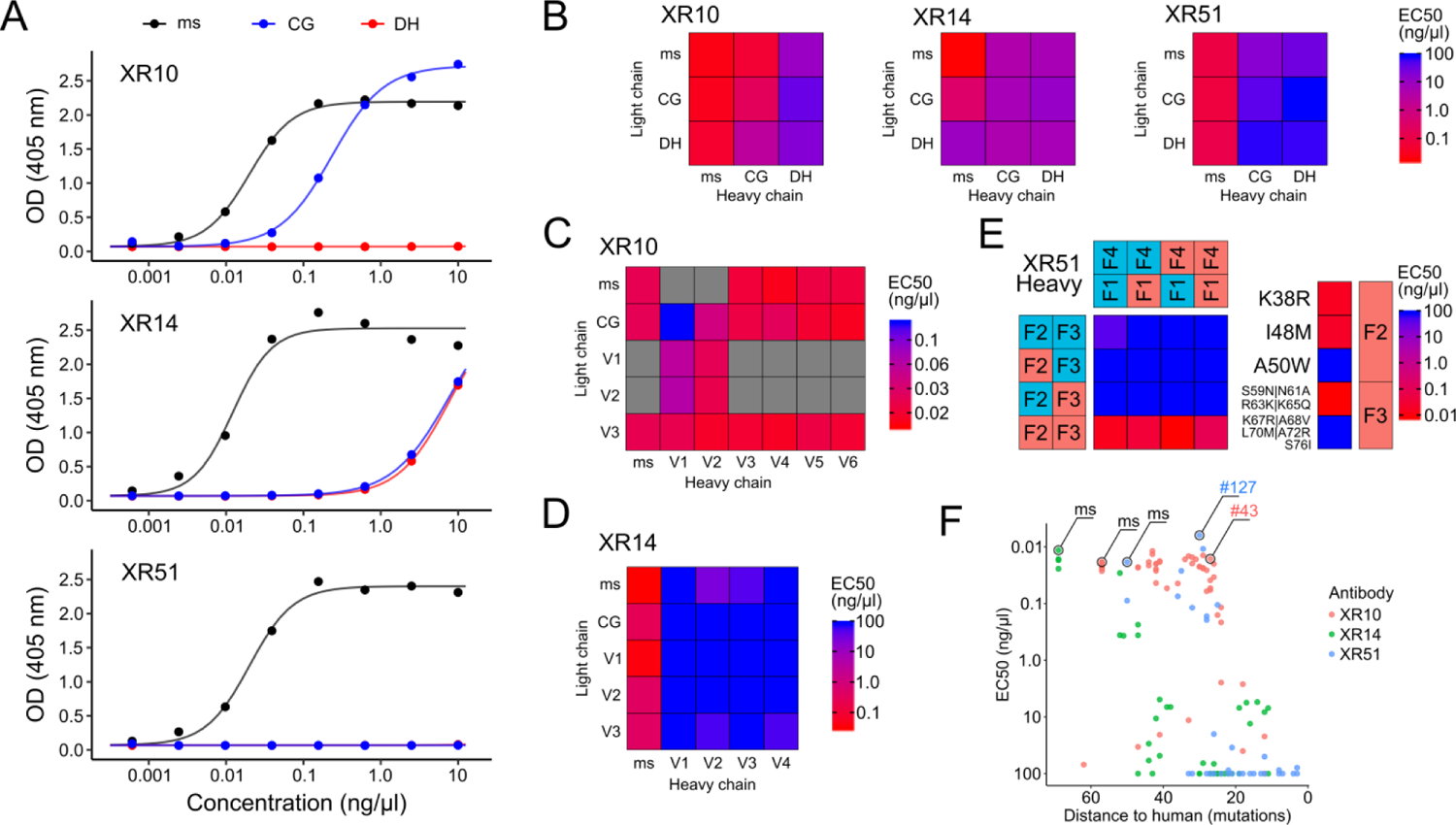
XR antibody humanization yielded highly humanized XR10 and XR51 variants with nominal binding affinity. **(A)** Mouse, CG, and DH variants binding. **(B)** Binding of XR antibodies generated by reshuffling of ms, CG, and DH heavy and light chains. **(C)** Binding of XR antibodies generated by framework reshuffling for XR10 **(C)** and XR14 **(D)** heavy and light chains, and for heavy chain of XR51 **(E)**. **(F)** Binding of XR variants plotted against the level of humanization. All binding assays performed with SARS-CoV-2 Wuhan RBD. OD - optical density, EC50 - effective dose 50%. CG - CDR-grafted version, DH - deeply humanized version.

### Different XR10 Vernier Zone residues contribute to Wuhan and Omicron BA.1 binding

In order to study how the humanization influences the binding of XR10 to different SARS-CoV-2 strains, we generated 110 antibody variants by reshuffling frameworks and chains of XR10 (Table S2, Figure S1) and tested their binding against Wuhan and Omicron BA.1 RBD (Figure 3A,B, Table S1). Only a minor fraction of the variants bound Omicron BA.1 RBD with affinity comparable to their anti-Wuhan interaction (Figure 3C, Figure S4A). DtH analysis revealed that Omicron BA.1 binding is extremely sensitive to humanization, with no variants of DtH≤35 binding with nominal affinity (Figure 3D). We hypothesized that Omicron BA.1 binding is influenced by a different set of AA exchanges introduced with the humanization, hence we used decision tree regression to determine the relative contribution of the mutated residues (Figure 3E). While humanizing H69, H102, and K37 residues (H – heavy, K – kappa) affect both Wuhan and Omicron BA.1 binding, H31 influences Wuhan binding exclusively and K58 and K84 contribute to Omicron BA.1 binding (Figure 3F). Mapping the residues on AlphaFold-generated msXR10 model demonstrated that K58 and K84 residues are distant from the paratope (Figure 3G) [45]. These results suggest a distinct mode of XR10-Omicron BA.1 interaction, where K58 and K84 AAs are critical for Omicron BA.1 binding proficient paratope folding.

**Figure 3.**
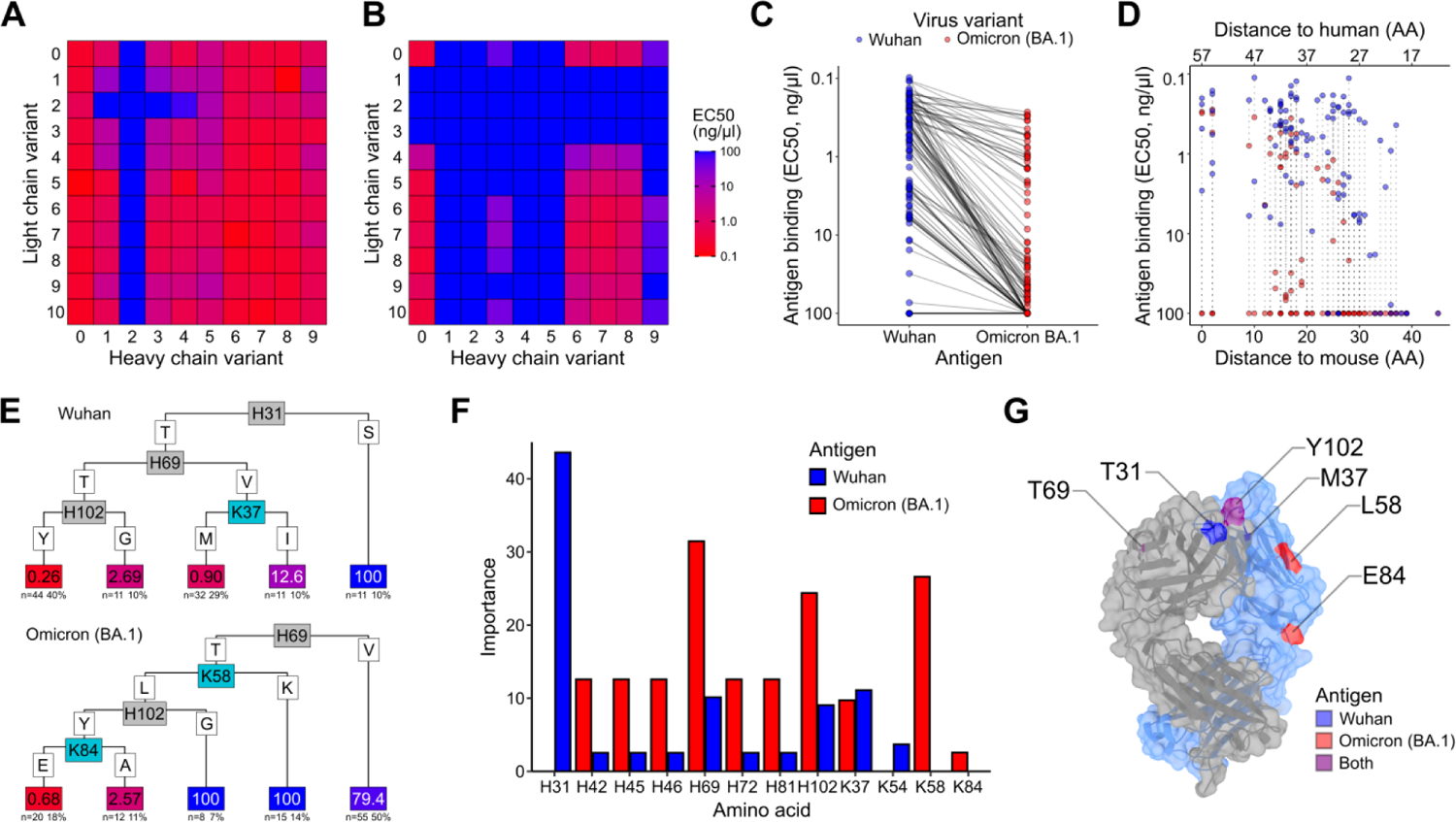
Different Vernier zone residues are contributing to XR10 binding to Wuhan and Omicron BA.1 SARS-CoV-2 RBD. All combinations of framework reshuffled XR10 heavy and light chains were expressed and profiled by binding to Wuhan **(A)** and Omicron BA.1 RBD **(B)**. **(C)** XR10 variants binding to Wuhan Omicron BA.1 RBD. **(D)** Binding to Wuhan and Omicron BA.1 RBD versus distance to human. **(E)** Decision tree regression analysis performed for Wuhan and Omicron BA.1 RBD binding **(F)** Contribution of individual Vernier Zones amino acid residues to RBD interaction. **(G)** Crucial amino acid residues mapped on the predicted mouse XR10 model.

### XR10v48 anti-SARS-CoV-2 mAb intranasal prophylactic protection in hamsters

In order to test how the humanization affected the neutralization potency of XR10 *in vivo*, we selected the most potent XR10 variants (EC_50_ < 1 ng/µl, XR10v51 as a low-affinity control) with DtH≤41 for further experiments (Figure S4A). We tested the selected variants in live virus neutralization for Wuhan, Delta, and Omicron BA.1 strains (Figure S4B). XR10v48 demonstrated the best combination of humanization extent (DtH=34, 85.3% homology to HGL), viral neutralization (IC_50_ of 3.0, 2.4, and 12.3 µg/ul for Wuhan, Delta, and Omicron BA.1, correspondingly) and higher affinity in comparison to mouse mAb (0.917 nM versus 1.167 nM for msXR10, Figure S5, Table S1). To evaluate XR10v48 performance in hamster protection, we administered 50 µg/kg of either humanized or mouse mAb intranasally simultaneously with SARS-CoV-2 Delta infection (4 animals per group, TCID 10^6^ per animal). Hamsters that received either mouse or humanized mAb, demonstrated a significant reduction of weight loss (Figure 4A,B). Although we did not detect significant decrease of SARS-CoV-2 load in nasal turbinates, lung viral load was significantly reduced both in mouse XR10 (Figure 4C) and XR10v48 administration (Figure 4D). We assessed the pulmonary damage by hemorrhagic foci counting (Figure 4E,F) and lung lesion score (see Methods, Figure 4G,H). For both the mouse XR10 and humanized XR10v48 treatment groups, lesions were less pronounced than in challenge-only controls (Figure 4E-H). These data suggest that XR10v48 can be used in SARS-CoV-2 infection prophylaxis at a relatively low single-administration dose. In conclusion, CRVZ humanization yielded an antibody variant with >80% homology to HGL, retained binding activity to SARS-CoV-2 spike, and *in vivo* protective activity.

**Figure 4.**
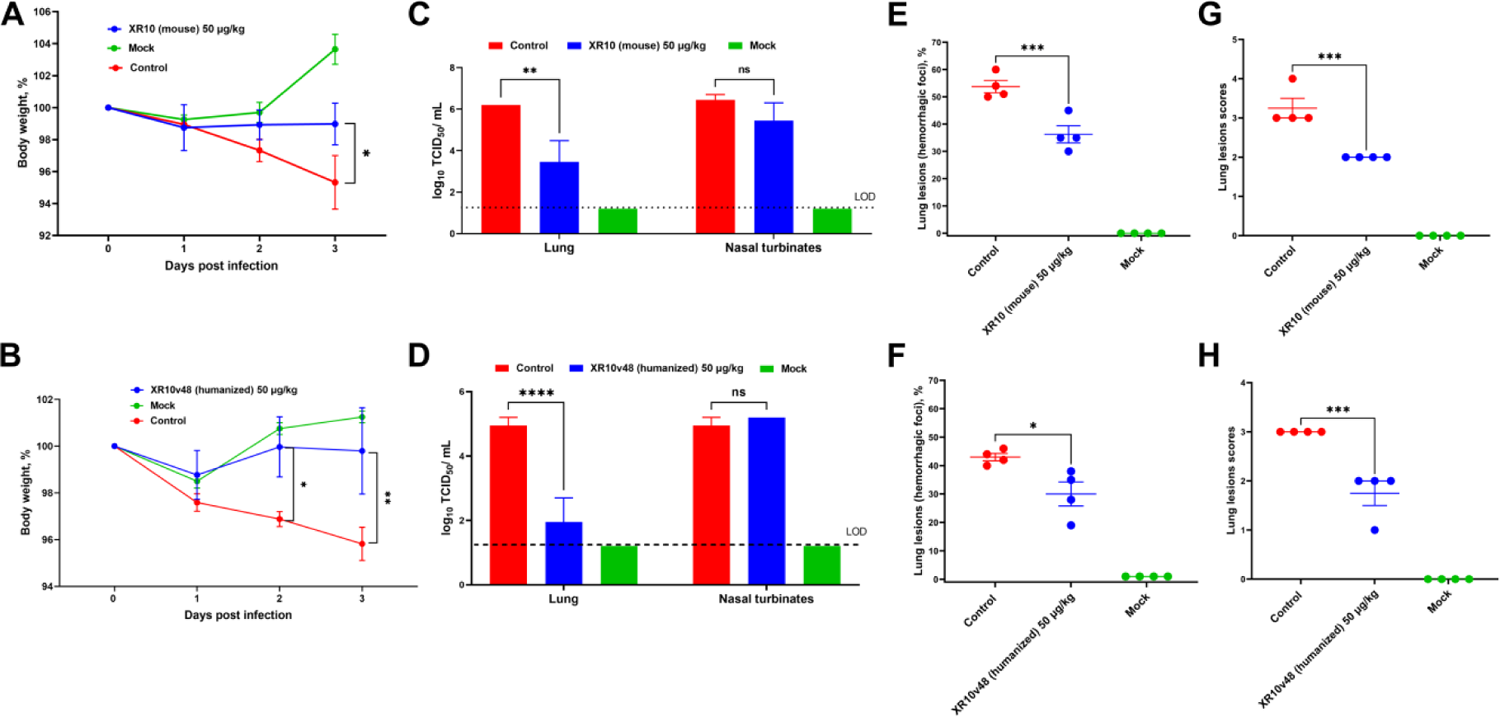
Intranasal prophylactic SARS-CoV-2 protection of mouse and humanized XR10 mAb in Syrian hamsters. Shown are changes in body weight at 0-3 days after challenge and treatment with mouse XR10 **(A)** or XR10v48 **(B)**; viral load in nasal turbinates and lungs expressed as log10 TCID50/mL on day 3 after challenge for mouse XR10 treatment **(C)** or XR10v48 administration **(D)**; lung lesions (hemorrhagic foci) **(E,F)** and lung histology analysis **(G,H)** on day 3 after challenge for mouse XR10 **(E,G)** or XR10v48 **(F,H)** administration. Differences between groups were assessed by Dunnett’s multiple comparison test. P<0.05 was considered significant. *P<0.05, **P<0.01 and ***P<0.001 and ****P<0.0001.

## DISCUSSION

Recombinant monoclonal antibodies represent promising candidates for therapy and prophylaxis of infectious diseases, as they are relatively stable and can be produced with high scalability at a low cost. Moreover, intranasal delivery of monoclonal antibodies was shown to confer viral protection in a prophylactic setting in humans and mice [46–48] and therapeutic setting in the hamster model [49].

In this study, we used hybridoma technology to generate a set of neutralizing antibodies with high affinity towards the SARS-CoV-2 Spike protein. These mAbs bound to various epitopes overlapping with the ACE2 interaction zone and demonstrated strong binding with affinity up to 1.089 nM. To decrease the probability of HAMA responses, we performed humanization of the generated antibodies. We used CRVZ humanization, a straightforward experiment-driven pipeline that allowed us to arrive at a high level of HGL homology (88.3% and 87.2% for XR10v43 and XR51v127) with nominal or higher affinity in comparison to mouse mAb. CRVZ can be further improved if combined with *in silico* humanization methods. For instance, recent advances in deep learning and language modeling combined with vast antibody repertoire databases yielded a platform able to humanize and evaluate the humanness of mouse mAbs, that suggested a similar set of amino acid exchanges for most of the sequences used in our study [44,50]. Additionally, molecular dynamics simulations can be used to predict the effect of humanization on the mouse mAb function [51]. These methods might aid the humanization and, on the other hand, can benefit from the binding, neutralization and *in vivo* protection data, acquired in the course of our study. Furthermore, humanization procedure might be circumvented by using genetically engineered mice [52] or phage display [53], however, these techniques might be technically complex and of limited availability.

It is important to note that, despite of widespread use of humanization procedure, the immunogenicity of some therapeutic mAbs remains unpredictable. Even fully human mAbs were shown to be immunogenic in clinical setting [54]. Although such methods as removing T-cell epitopes (de-immunization [55]) or introducing T-reg epitopes (tolerization [56,57]) were successfully applied to mitigate the anti-therapeutic response, further research is needed to control the immunogenicity of mAbs.

The humanized XR10v48 (85.3% homology to HGL) selected for *in vivo* testing was as efficient in protection as the original mouse variant. We showed protection against infection with SARS-CoV-2 Delta virus in Syrian hamster model at a single-dose nasal administration at a concentration of 50 ug/kg, which is substantially lower than previously reported nasal therapeutic antibodies [49]. The delivery route chosen for XR10v48 offers a significant advantage for treatment and prophylaxis of respiratory viruses. Even though it is possible to prevent infection with the high-dose intravenous mAb administration, the systemic administration was shown to result in a poor distribution into the lung [58,59]. Although we did not analyze the biodistribution of the mAb in the mice, we show the significant reduction of the viral load in the lung tissues, which suggests its successful delivery to the primary site of infection. Intranasal administration of IgG was previously shown to lead to FcRn-mediated uptake through the mucosa [60]. Albeit this transfer was suggested to contribute to mAb degradation, it also enhances the processing by the antigen-presenting cells [61,62]. These findings highlight the potential utility of intranasally delivered mAb and prompt further research in this direction.

In conclusion, the study presents a straightforward humanization pipeline that requires no specialist knowledge and led to the development of a highly humanized antibody. Its product, XR10v48, binds to SARS-CoV-2 Spike with subnanomolar activity and can be used for prophylaxis of SARS-CoV-2 infection. XR10v48 is delivered intranasally providing a non-invasive alternative to injected therapeutic antibodies. Because a single-dose administration and low concentration are required for antiviral protection, XR10v48 may reduce the economic burden in a clinical application.

## MATERIALS AND METHODS

### Mouse immunization

Four-to-six-week-old female BALB/c mice were immunized 4 times every three weeks with 50 ug of the Wuhan Spike or RBD protein via intraperitoneal injection with complete Freund’s adjuvant. After 4 doses of the same injection, the response potency was estimated using antigen ELISA. Booster dose was injected for high responders 20 days after the fourth immunization. 4 days after the boost, mice were sacrificed and spleens were harvested.

### Hybridoma generation

Splenocytes of the mice were harvested and fused with mouse myeloma SP2.0 cells. Fused cells were cultured in DMEM supplemented with 10% FBS, HAT medium, hybridoma growth factors in 96-well culture plates. 2 weeks after fusion, the supernatants were analyzed for antibody titer with ELISA. Selected clones were subcloned via limiting dilution.

## ELISA

In-house-produced recombinant RBD was immobilized on a high-binding 96 well ELISA plate (Corning, #CLS3690) at 4 µg/ml in PBS (Sigma Aldrich, MO, USA) overnight at +4°C. Plates were blocked for 1h with 1% BSA (Thermo Fisher, Gibco, MA, USA) in PBS at room temperature. Antibodies were diluted in PBS 1% BSA to indicated serial dilutions, added to coated plates and incubated for 1h at room temperature. Plates were developed with an anti-human IgG-alkaline phosphatase (AP)-coupled antibody (SouthernBiotech #2040-04) diluted 1:500 in PBS 1% BSA. Bicarbonate buffer with 4-Nitrophenyl phosphate disodium salt hexahydrate substrate (Sigma, #S0942-50TAB) was added and absorbance was measured at 405 nm in a Cytation 5 device (BioTek). Between all indicated incubation steps, plates were washed 3 times with PBS 0.05% Tween-20. 50% of maximum IgG binding to RBD or nucleocapsid (ED50) was determined for each sample by sigmoid curve fitting with non-linear regression performed in R (stats package) [63]. For curve fitting, upper and lower plateaus of the reference antibody (murine antibodies) were applied to all samples. The positive control was used to normalize independent measurements.

### Competition ELISA

Upon 1 h blocking with PBS/1% BSA, serially diluted antibodies in PBS/1% BSA were added to SARS-CoV-2 RBD-coated (10 µg/ml) 96 well plates. After 1 h of incubation at room temperature, self-produced biotinylated human ACE2-hFcg1, EY6A, P2B-2F6, P2C-1F11, REGN10933, REGN10987, CC12.1, C144, S2H13, S309, 4A8 were added to a 50% final effective concentration (EC_50_) in PBS/1% BSA for competition with monoclonal antibodies. After another hour at room temperature, plates were incubated with an AP-coupled streptavidin (Southern Biotech, #SBA-7105-04) at a 1:500 dilution in PBS/1% BSA to detect biotinylated ACE2-hFcg1 that was not prevented by serum antibodies from binding to RBD. The 50% blocking dose (BD_50_) was determined by sigmoid curve fitting with non-linear regression performed in R (stats package). Upper and lower plateaus of the (non-)biotinylated ACE2-hFcg1 control served as a reference.

### Monoclonal antibody sequencing

Sequencing of XR10, XR14, and XR51 antibodies was performed by Absolute Antibody Ltd (UK) by whole transcriptome shotgun sequencing (RNA-Seq).

### Recombinant proteins cloning

SARS-CoV-2 RBD WT was cloned containing the signal peptide spanning amino acids M1-Q14 and R319-F541 of the pCAGGS-SARS-CoV-2 RBD Wuhan plasmid (kindly provided by F. Krammer; GenBank: MN908947.3, [64]), followed by a C-terminal Twin-Strep-tag sequence (WSHPQFEKGGGSGGGSGGSAWSHPQFEK) and a hexahistidine tag. SARS-CoV-2 spike encoding plasmid comprised the amino acids M1-Q1208 of the Wuhan SARS-CoV-2 variant (GenBank: MN908947.3) with Furin cleavage site mutations (682-685 RRAR -> QQAQ) as well as 6 Proline mutations (HexaPro) to generate a protease-resistant, stable and highly immunogenic SARS-CoV-2 spike protein locked in the pre-fusion conformation followed by a Twin-Strep-tag and a hexahistidine tag. For expression of the membrane-bound version, a SARS-CoV-2 spike full-length vector with a C-terminal 19 amino acid deletion comprising amino acids M1-C1254 of the Wuhan SARS-CoV-2 variant (GenBank: MN908947.3) was used. Mutations of RBD and full spike were introduced by PCR mutagenesis. The N-terminal domain (amino acids M1-S305) of the SARS-CoV-2 spike was cloned to a C-terminal Twin-Strep-tag and a hexahistidine tag. Plasmids containing human full-length ACE2 (Clone: OHu20260C; M1-F805) and rabbit full-length ACE2 (Clone: OOb21562C; M1-F805) C-terminally linked to enhanced green fluorescent protein (eGFP) were purchased from Genscript (GenScript Biotech Netherlands B.V.). Heavy and light chain sequences of humanized and control mAbs were synthesized by oligonucleotide assembly and cloned using Gibson assembly (NEBuilder HiFi DNA assembly kit, NEB, #E5520S) into IgG1 heavy and kappa or lambda light chain expression vectors from Oxgene (Sigma-Aldrich; #PP2409-1KT): class 1-4 anti-SARS-CoV-2 RBD: EY6A, P2B-2F6 and P2C-1F11, REGN10933 and REGN10987, CC12.1, C144, S2H13, S309, anti-SARS-CoV-2 NTD: 4A8. Framework reshuffling for humanized mAbs was performed using framework and CDRs amplification and subsequent Gibson assembly. Human ACE2-hFcg1 was produced by fusing recombinant human ACE2 Q18-V739 fragment to human IgG1-Fc (E99-K330 portion, where the first amino acid is G encoded by J-CH1 fusion).

### HEK cell recombinant protein production

Cloning constructs were used to transfect FreeStyle 293-F cells that were grown in suspension using FreeStyle 293 expression medium (Life Technologies, #A1435101) at 37°C in a humidified 8% CO2 incubator rotating at 125 rpm. Cells were grown to a density of 2.5 million cells per ml, transfected using polyethyleneimine (PEI, Polysciences Europe GmbH; #23966-1; 4 µg/ml in cell suspension) and DNA (1200 ng/ml in cell suspension), and cultivated for 3 days. The supernatants were harvested and proteins purified by His SpinTrap columns (for His-tagged antigens) or Protein G columns (for the antibodies) according to the manufacturer’s instructions (Cytiva, His: #28-9321-71, Ab: #28-4083-47). The eluted protein was transferred to phosphate-buffered saline (PBS) via buffer exchange using Amicon Ultra-4 ultrafiltration column with 50 kDa or 10 kDa cutoff (Millipore, #UFC805008, #UFC801096). Protein concentration was determined by His-tag specific ELISA using a mouse anti-His-tag antibody (Abcam, #ab18184) and a goat anti-mouse IgG Fc antibody conjugated to alkaline phosphatase (Southern Biotech, #SBA-1033-04) as a detection reagent. Antibody concentration was determined by absorbance measurement at 280 nm. Protein production was confirmed by SDS-PAGE and western blot using a mouse anti-His antibody (Abcam, #ab18184) and an IRDye 800CW donkey anti-mouse antibody (Li-Cor Biosciences, #925-32212).

### Insect cell recombinant spike protein production

The antigen in SpikoGen® vaccine corresponds to aa 14 – 1213 of the spike protein sequence of the ancestral Wuhan-Hu-1 strain (accession number: NC 045512) with various modifications, as previously described [18]. In this study, Wuhan, Beta and Gamma, Delta and Omicron spike proteins were manufactured using the same process as the human approved SpikoGen® vaccine. Spike antigens and Advax-CpG55.2 adjuvant were provided by Vaxine Pty Ltd, Adelaide, Australia.

### Administration of monoclonal antibody to hamsters and their SARS-CoV-2 virus challenge

Each of the monoclonal antibody preparations XR10 (1.24 mg/mL) and XR10RH (4.73 mg/mL) were diluted with DMEM at a final concentration of 5.5 µg/100 µL or 50 µg/kg, and antibodies or PBS intranasally administered to 6-8-week-old male Syrian hamsters (50 µL in each hamster nostril, n=4/group) under intraperitoneal ketamine (100 mg/kg) and xylazine (10 mg/kg) anesthesia (Table 1). Syrian hamsters were obtained from the laboratory animal nursery of the M. Aikimbayev National Scientific Center for Especially Dangerous Infections (NSCEDI, Kazakhstan) and during the experiment housed an individually ventilated system with a 30-cage ventilated rack in the ABSL-3 laboratory of NSCEDI. Two hours later, the hamsters of groups 1-3 under anesthesia were intranasally infected with the strain hCoV-19/Kazakhstan/KazNARU-NSCEDI-5526/2021 of Delta variant SARS-CoV-2 virus (full protein sequence was published in GISAID database under number EPI_ISL_4198501) at a dose of 10^6.0^ TCID_50_/100 µL (50 µL per nostril) as previously described [65]. Animals of the negative control group, which were kept in a separate room of the ABSL-3 laboratory from the experimental groups, were subjected to a similar anesthesia procedure. Infected animals were clinically monitored from day 0 to day 3, and their weight was measured daily. On day 3 after the challenge, animals from each group were sacrificed, and samples from nasal turbinates and lungs were collected. The lungs of the animals were evaluated for the level of lesions by the area of hemorrhage in the JMicroVision program (v 1.3.4). The percentage ratio of the area of the affected areas to the total area of the lung surface was studied. Two lobes of the left lung were homogenized in 1 ml of DMEM using a TissueLyser II (QIAGEN) device at 300 vibrations per minute for 60 s. The supernatant obtained after centrifugation (5000 g for 15 min at 4°C) was stored at −70°C for the determination of the virus titer. Three right lung lobes of each animal were fixed in 10% formalin for histopathological examination to confirm detected lung lesions.

**Table 1.**
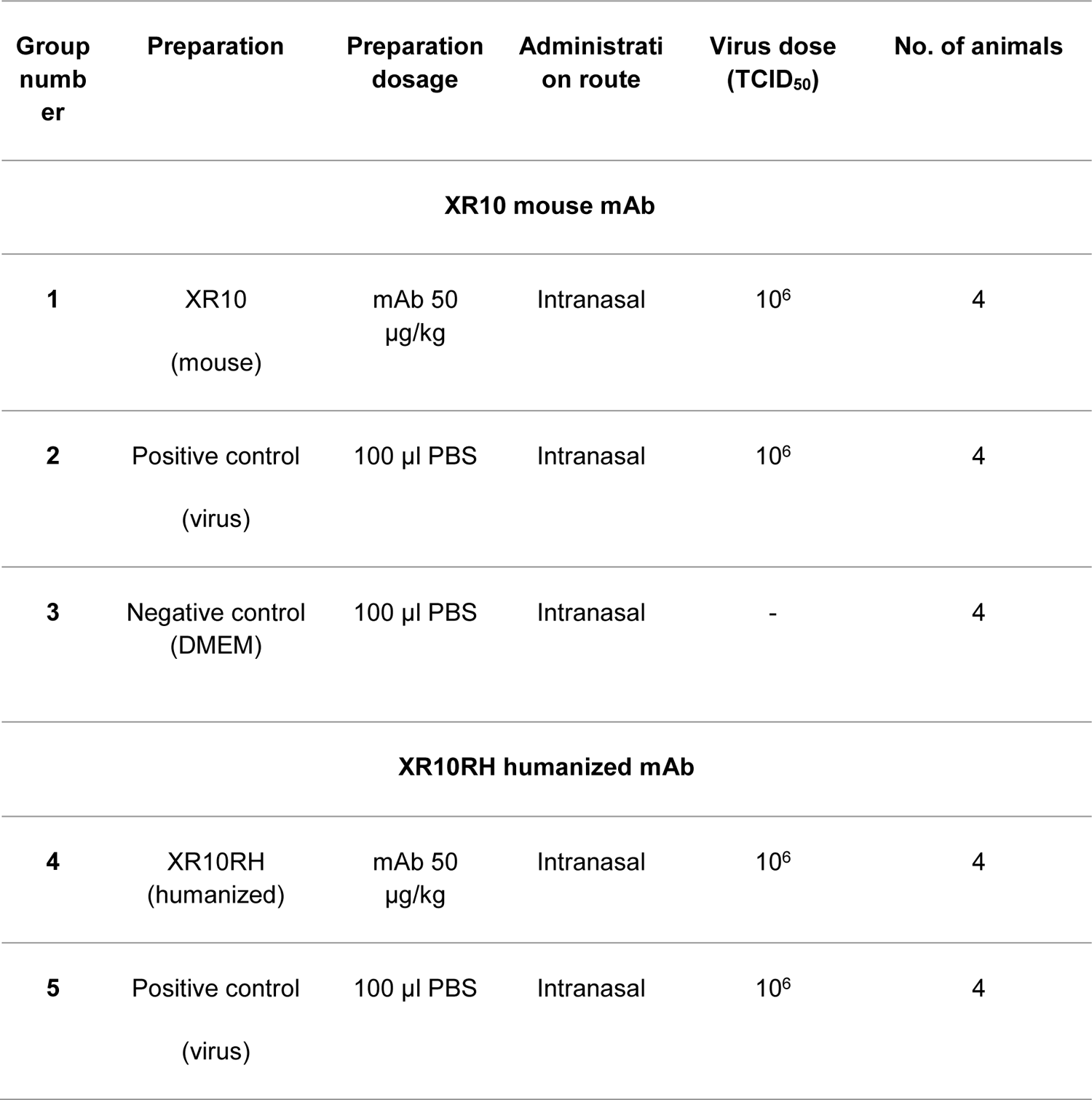
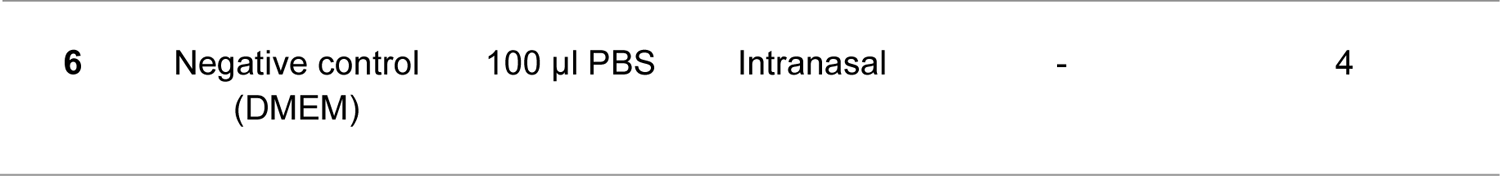
Study Design.

| Group number | Preparation | Preparation dosage | Administration route | Virus dose (TCID<sub>50</sub>) | No. of animals |
| --- | --- | --- | --- | --- | --- |
| **XR10 mouse mAb** |  |  |  |  |  |
| **1** | XR10<br>(mouse) | mAb 50<br>μg/kg | Intranasal | $10^6$ | 4 |
| **2** | Positive control<br>(virus) | 100 μl PBS | Intranasal | $10^6$ | 4 |
| **3** | Negative control<br>(DMEM) | 100 μl PBS | Intranasal | - | 4 |
| **XR10RH humanized mAb** |  |  |  |  |  |
| **4** | XR10RH<br>(humanized) | mAb 50<br>μg/kg | Intranasal | $10^6$ | 4 |
| **5** | Positive control<br>(virus) | 100 μl PBS | Intranasal | $10^6$ | 4 |
| 6 | Negative control<br>(DMEM) | 100 µl PBS | Intranasal | - | 4 |

### Determination of the infectious virus load

Virus titers in homogenates of respiratory tract tissues were determined using the 50% tissue culture infectious dose (TCID_50_) analysis. The tissue homogenates were diluted ten-fold in a medium (DMEM-2% FCS-1% antibiotic-antimycotic) and transferred in quadruplicate into 96-well plates containing Vero-E6 confluent cells. The plates were then incubated at 37°C and 5% CO_2_ for 5 days. The titration results were visually calculated by studying the cellular monolayer under a microscope for specific cytopathic effects such as cell rounding and cell separation from the monolayer. The viral titer was determined using the Reed and Muench method and expressed as log_10_ TCID_50_/mL.

### Histological analysis of hamster lungs

The lungs of hamsters were fixed in 10% neutral buffered formaldehyde after excision, washed with water, and treated with four portions of 100% isopropyl alcohol and two portions of xylene. The material was then soaked in four portions of paraffin, and blocks were created. Histological blocks were sectioned to a thickness of 5 µm using a microprocessor microtome MZP-01 (CB Technom, Russia). The sections were dewaxed in two portions of xylene and three portions of ethyl alcohol with reduced concentrations (96%, 80%, 70%), then stained with hematoxylin (#05-002, BioVitrum, Russia) and eosin (#C0362, DiaPath, Italy). Subsequently, the sections were clarified in increasing concentrations of ethyl alcohol (70%, 80%, 96%) and two portions of xylene. The sections were covered with glass coverslips using the synthetic medium Bio Mount (#2813, Bio Optica, Italy). The slides were examined using the Mshot microscope (China), model MF52-N. Photos were taken with a magnification of x40 using Mshot MS23 camera tips (China) and the MShot Image Analysis System program (China). Microscopic examination of the lungs followed classical canons adopted for parenchymal organs, and the description of pathological conditions induced by SARS-CoV and SARS-CoV-2 was used for narration. Each slide was quantitatively assessed based on the severity of histological changes, including interstitial pneumonia, alveolitis, bronchiolitis, alveolar destruction, interstitial infiltration, pulmonary hemorrhage, and peribronchiolar inflammation. For evaluation of pathological changes in peribronchiolar and perivascular infiltrates, a scoring system was used: 4 points - extremely severe; 3 points - severe; 2 points - moderate; 1 point - mild; 0 point - no changes.

### Biosafety and bioethics

All operations involving SARS-CoV-2 and viral experiments on animals were conducted in the BSL-3 and ABSL-3 laboratories of NSCEDI, where the international standard ISO 35001:2019 “Biological Risk Management for Laboratories and Other Related Organizations” was followed. Laboratory animals were housed in individually ventilated cages (Techniplast, Italy, and Allentown, USA) under a 12/12 day and night cycle. This study was conducted in compliance with national and international laws and guidelines for the handling of laboratory animals. The protocol was approved by the Institutional Animal Care and Use Committee of the National Scientific Center of Especially Dangerous Infections, Protocol No. 4, dated September 22, 2020.

### Statistical analysis

The statistical data analysis was performed using the GraphPad Prism program, version 9.0.0 (San Diego, California, USA) and R, version 4.3.1 [63]. Differences in weight loss, viral load, and pathological changes in the lungs between the groups of animals were assessed using the Tukey multiple comparison test. A Shapiro-Wilk test was used for normality testing of continuous variables. An independent *t*-test was used when continuous data met the criteria of the normality test. Otherwise, the Mann-Whitney U test was used. Spearman correlations were performed with a 95% confidence interval. The limit of detection for viral titer was 0.7 log_10_ TCID_50_/0.2 mL. A p-value of less than 0.05 was considered statistically significant for all comparisons. Figures were generated using R ggplot package, Inkscape software, and BioRender.

### Protein structure analysis

Antibody structures were predicted using templated AlphaFold-2 software [45]. Antibody-antigen interaction was analyzed using the HDOCK server. Protein structures and interactions were analyzed and depicted with UCSF ChimeraX 1.173. The following PDB codes were used: 6M0J for SARS-CoV-2 RBD and ACE2 [4], 6ZER for EY6A [34], 6XC3 for CC12.1 [66], 8DCC for P2B-2F6 [67], 8GX9 for P2C-1F11 [68], 6XDG for REGN10933 and REGN 10987 [33], 7K90 for C144 [69], 7JV2 for S2H13 [40], 7XSW for S309 [70], 8DM4 for 4A8 [71].

## ACKNOWLEDGEMENTS

N.P. was supported by the funding from National Institute of Allergy and Infectious Diseases of the National Institutes of Health under Contracts HHS-N272201400053C and HHS-N272200800039C. The content is solely the responsibility of the authors and does not necessarily represent the official views of the National Institutes of Health. We would like to thank Dr. Mariia Sergeeva from Smorodintsev Research Institute of Influenza for performing pseudovirus neutralization assay. We extend our gratitude to Turebekov N. and Fomin G. for their diligent efforts in ensuring the biosecurity and safety aspects of our research, as well as for conducting the histological studies on hamster lung tissue samples. We also thank Zhambyrbayeva L.S. and Sarmantayeva K.B. for their exceptional care and maintenance of the hamsters.

## SUPPLEMENTARY FIGURES

**Figure S1.**
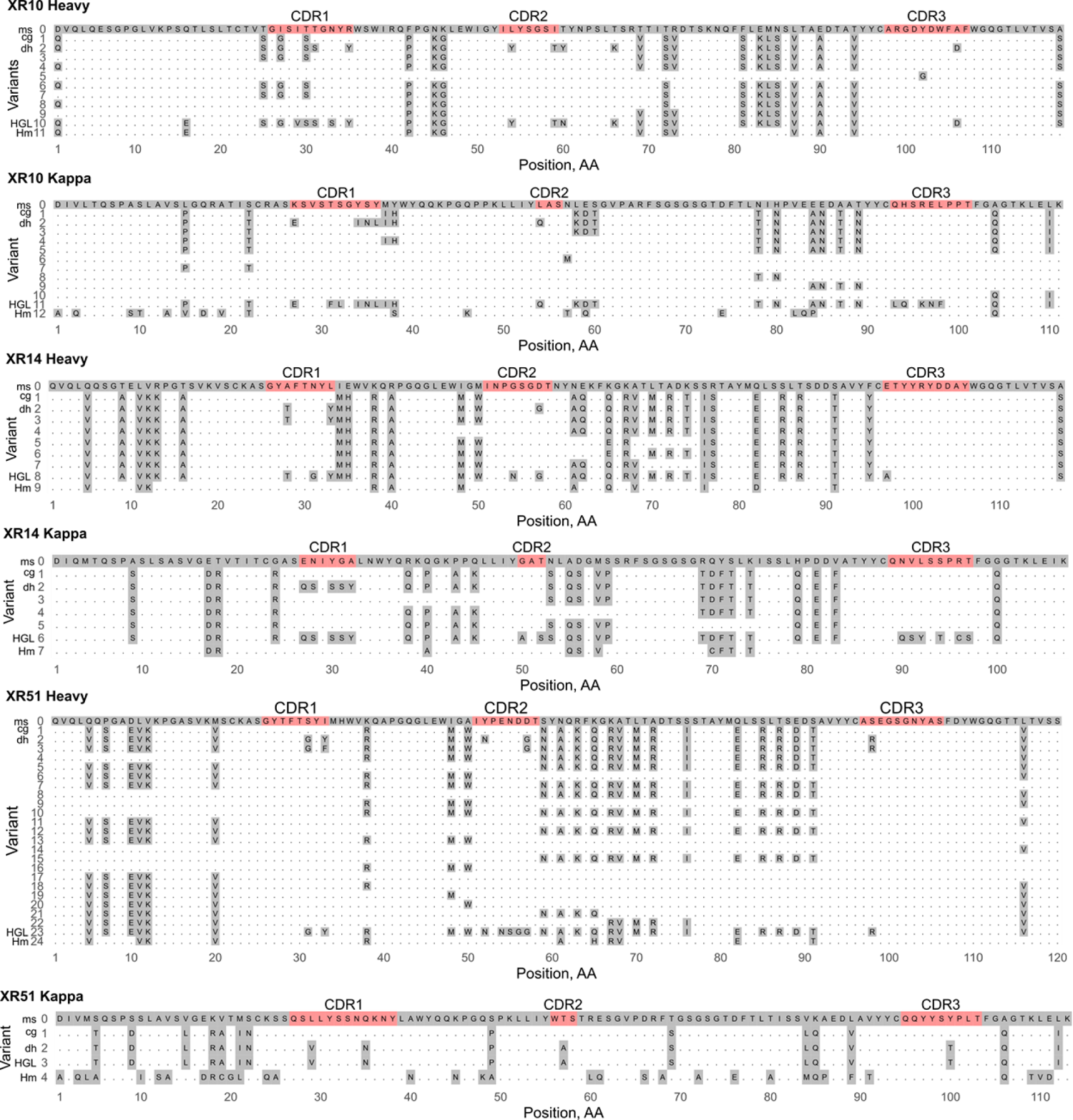
Multiple alignment of the XR amino acid sequences. ms – original mouse mAb sequence, cg – CDR grafted version, dh – deeply humanized version, HGL – germline human sequence, Hm - sequence humanized by Hu-mAbs.

**Figure S2.**
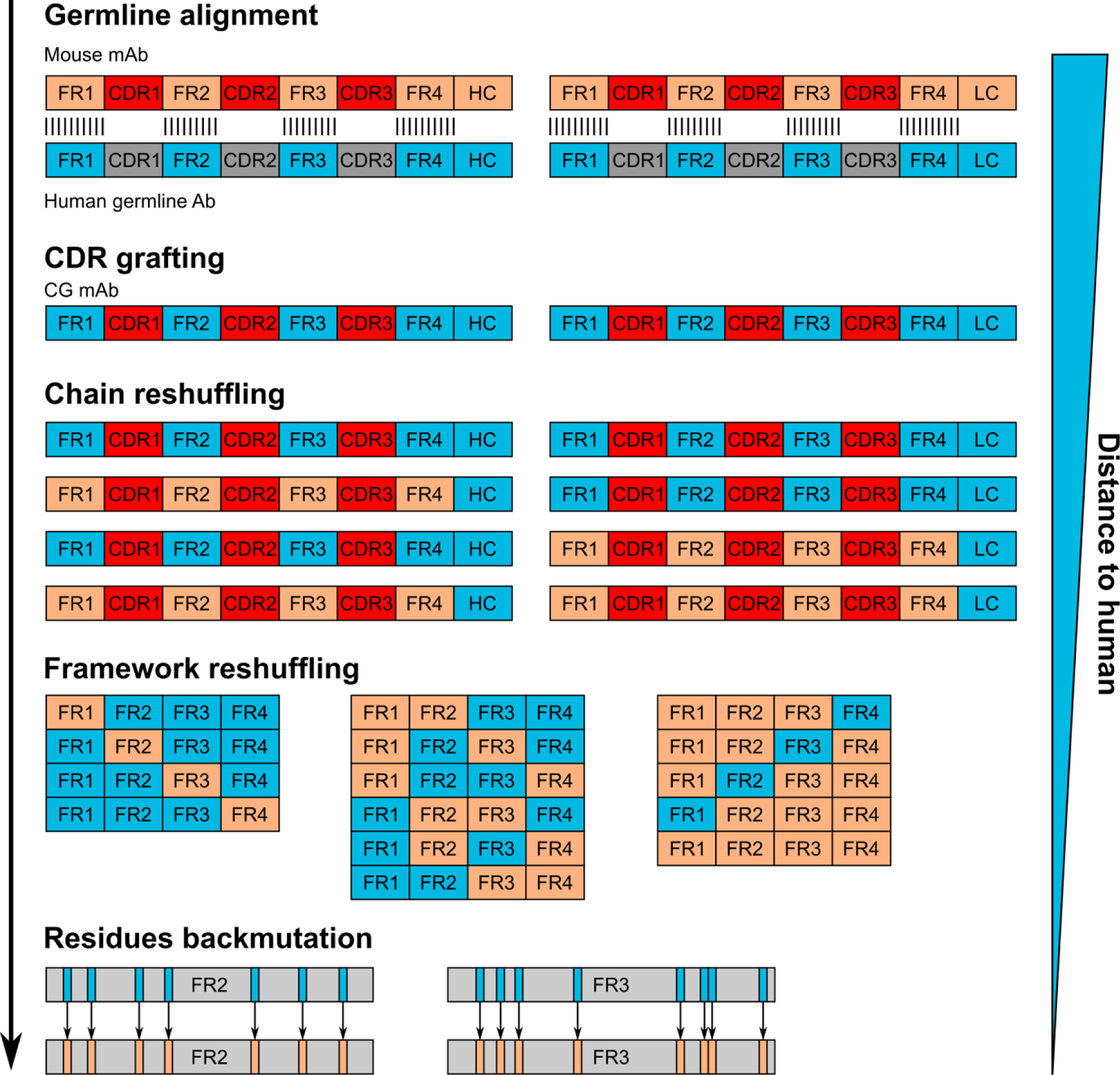
Experimentally-driven mAb humanization procedure outline. The first step entails alignment to human (or target organism) germline segment library to select the framework regions of the highest homology. CDRs are then grafted in the human frameworks. Chain and framework reshuffling is used to locate the binding-affecting mutations introduced through humanization. Final step is performed on individual amino acid residues level.

**Figure S3.**
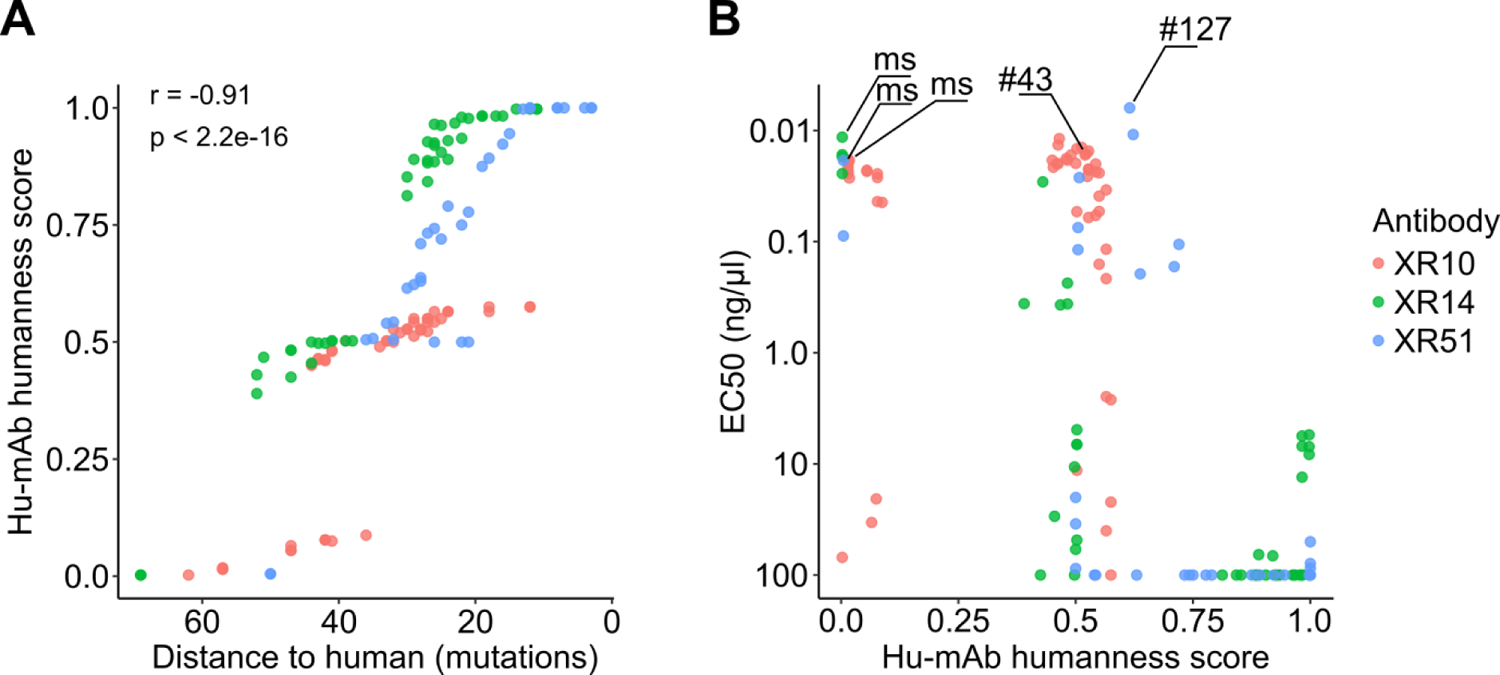
Variants evaluation by Hu-mAbs software. **(A)** Humanization extent estimated by Hu-mAb humanness score and DtH. Correlation is calculated with Spearman’s rank test. **(B)** Antibody binding (EC50) to Wuhan RBD against Hu-mAb humanness score. ms - original mouse mAb variant.

**Figure S4.**
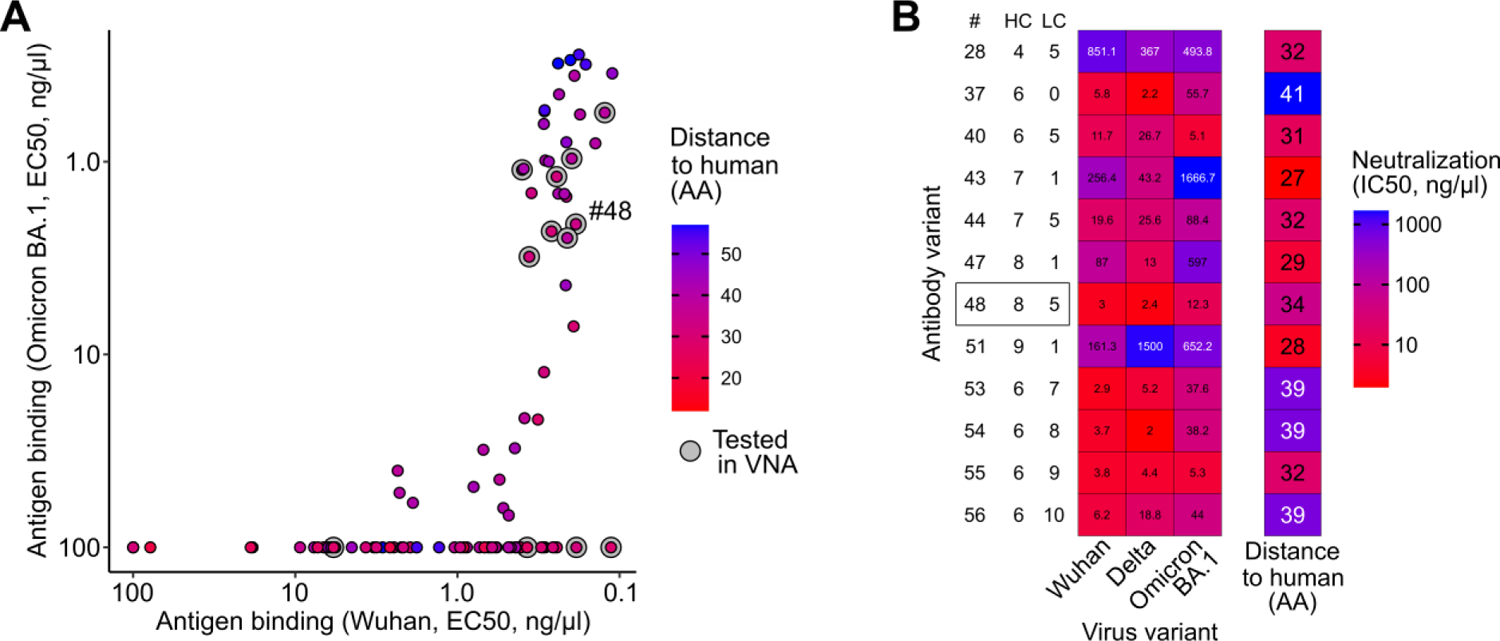
Subset of XR10 variants characterization. **(A)** Binding of all XR10 variants to Wuhan and Omicron BA.1 RBD with the further tested subset highlighted. **(B)** Neutralization of Wuhan, Delta, and Omicron BA.1 viruses by the selected XR10 antibodies. XR10v48 is selected for hamster challenge tests.

**Figure S5.**
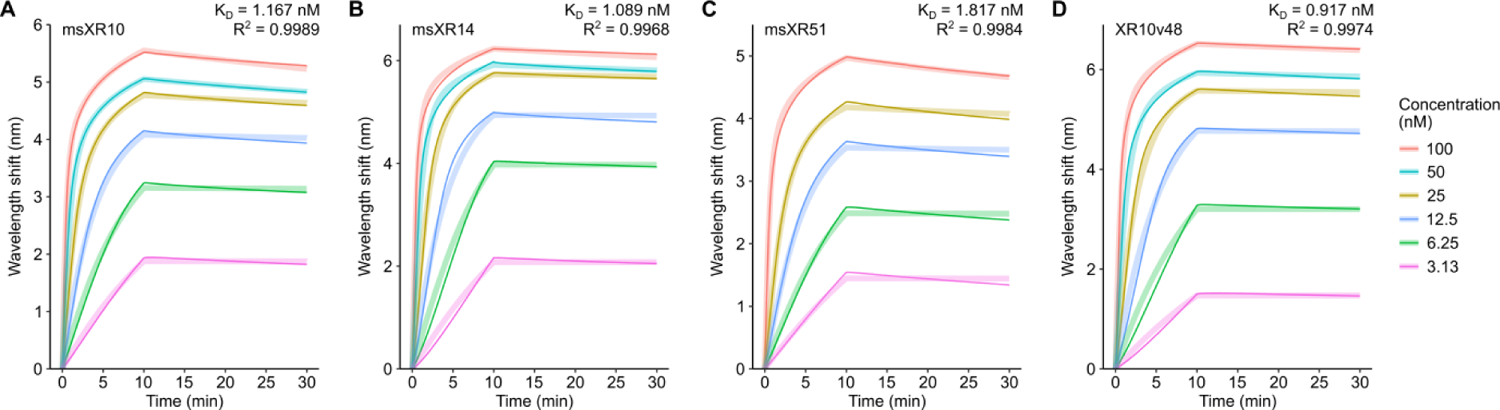
Biolayer interferometry analysis. Kd calculated using the global fit of the association/dissociation curves received from wavelength shift measurement. R^2^ – determination coefficient.

